# Sex-dependent endocannabinoid adaptations in supraspinal pain-modulatory circuits after peripheral nerve injury

**DOI:** 10.64898/2026.10.15.759453

**Authors:** Laura Boullon, David P. Finn, Álvaro Llorente-Berzal

## Abstract

Peripheral nerve injury produces persistent pain-related behaviours that differ between sexes, but whether chronic neuropathic pain is accompanied by sex-dependent alterations in endocannabinoid signalling across pain-modulatory circuits remains unclear. We quantified endocannabinoids and related *N*-acylethanolamines and expression of cannabinoid receptors and metabolic enzymes across supraspinal and spinal regions 100 days after spared nerve injury (SNI) in male and female rats. SNI increased 2-arachidonoylglycerol (2-AG) and palmitoylethanolamide (PEA) in the prefrontal cortex and 2-AG in the rostral ventromedial medulla in males, with exploratory associations between these measures and reduced pain-related hypersensitivity. In females, SNI increased monoacylglycerol lipase expression in the periaqueductal grey. Additional sex- and hemisphere-dependent differences in ECS components were observed in limbic regions, whereas no significant injury-related changes were detected in spinal cord or plasma. These findings identify regionally distinct ECS alterations during established neuropathic pain and suggest that chronic nerve injury is associated with different patterns of supraspinal ECS plasticity in males and females.

**Perspective:** This study identifies sex- and region-dependent alterations in endocannabinoid signalling during established neuropathic pain. Distinct changes in supraspinal pain-modulatory regions in male and female rats highlight biological sex as an important consideration when investigating the endocannabinoid system as a therapeutic target for chronic neuropathic pain.

## Introduction

Despite extensive research highlighting the analgesic potential of the endocannabinoid system (ECS) in chronic pain disorders ^14,20,33,41,45^, a critical gap remains in understanding whether and how the ECS is modulated by chronic pain in a sex-dependent manner. Most studies have focused on male rodents, limiting our insight into sex-dependent ECS adaptations. Yet, evidence indicates that cannabinoid-mediated antinociception differs between sexes, with ECS-modulating compounds often showing greater antinociceptive efficacy in females than in males ^7,9,10,42,48,50^. Moreover, the mechanisms underlying these effects can differ by sex ^10^, suggesting that biological sex may influence the interaction between ECS signalling and pain modulation.

Pain etiology may further influence these sex differences. For example, delta-9-tetrahydrocannabinol (Δ^9^-THC) greater efficacy in attenuating Complete Freund’s Adjuvant-induced inflammatory pain in females than in males ^8^, but greater efficacy in males after sciatic nerve injury ^29^ Thus, different pain states may be accompanied by distinct ECS adaptations in males and females, potentially contributing to variation in cannabinoid-mediated antinociception. Although several studies have described ECS alterations after nerve injury in male rodents ^37,40^, considerably less is known about corresponding adaptations in females. Determining whether peripheral nerve injury alters ECS signalling differently in males and females is therefore important for understanding sex-dependent endocannabinoid regulation of persistent pain.

The descending pain-modulatory system, which includes key regions such as the periaqueductal grey (PAG), rostral ventromedial medulla (RVM), and spinal dorsal horn, is a key locus for ECS modulation of nociceptive transmission ^5,39^. Endocannabinoid signalling within this circuitry fine-tunes the balance between inhibitory and facilitatory control of pain, largely through CB_1_ receptor-mediated regulation of GABAergic and glutamatergic neurotransmission ^39^. Alterations in ECS signalling within these regions may therefore influence descending modulation of nociceptive processing. In addition to descending pain-modulatory regions, the prefrontal cortex, amygdala, and hippocampus contribute to the affective and cognitive dimensions of pain, with endocannabinoid signalling within these regions regulating stress responses, pain processing, and affective behaviour ^15,18,21,22,47^. Peripheral nerve injury may therefore be accompanied by regionally distinct ECS alterations across interconnected systems involved in sensory and affective dimensions of persistent pain.

The present study tested the hypothesis that peripheral nerve injury is associated with sex-dependent alterations in ECS components across pain-processing regions. Because ECS adaptations following nerve injury vary with pain state and time after injury ^37,40^, and there is a recognized need to examine the neurobiology of pain over longer post-injury periods ^38^, we characterized the ECS 100 days after spared nerve injury (SNI), when persistent pain-related behaviours are well established in both sexes ^3^. We quantified endocannabinoids and related *N*-acylethanolamines, together with cannabinoid receptors and principal catabolic enzymes, across cortical, limbic, descending pain-modulatory, and spinal regions in male and female rats. We further examined whether injury-associated ECS alterations were associated with mechanical and cold hypersensitivity. This approach was designed to determine whether established neuropathic pain is accompanied by distinct regional patterns of ECS alteration in males and females.

## Materials and methods

### Animals

Forty-eight adult (8-9 weeks old) male (∼250g) and female (∼200g) Sprague-Dawley rats (Charles River, Margate, United Kingdom) were used in the study, with a subset of twenty-four rats (n=6 per sex/group) randomly selected for the molecular assays reported here. To ensure uniform baseline sensitivity, rats were assigned to experimental groups using constrained randomization based on their baseline von Frey (vF) thresholds. The individual rat served as an experimental unit. Rats were excluded from statistical analysis if they were identified as outliers, defined as values exceeding ± 2 standard deviations from the group mean. Investigators were blinded to group allocation during both behavioral testing and the subsequent molecular procedures. Rats were pair-housed (sex- and surgery-matched) with water and food (14% Harlan Teklad 2014 Maintenance Diet, Envigo, Huntingdon, Cambridgeshire, United Kingdom) available *ad libitum* under a 12:12h light/dark cycle. All procedures, including Spared Nerve Injury or Sham surgeries, were performed during the light phase and were approved by the Animal Care and Research Ethics Committee, University of Galway, under license from the Health Products Regulatory Authority in the Republic of Ireland (per EU Directive 2010/63) ^25^. These procedures are reported in accordance with the ARRIVE 2.0 guidelines^12^.

### Experimental design

Brain and spinal cord samples from male and female Sprague-Dawley rats were collected to investigate potential alterations in the ECS 100 days following Spared Nerve Injury (SNI). An overview of the behavioural characterization of the model and the experimental procedures included in this study is provided in Figure 1, and the behavioural results have been reported in Boullon et al., 2025^3^. Pain-related behavioural data collected 24 hours before euthanasia were used for correlation analyses with molecular and biochemical measures, for the focus of the present paper. Endocannabinoid ligands - anandamide (AEA) and 2-arachidonoylglycerol (2-AG) - and related *N*-acylethanolamines - palmitoylethanolamide (PEA) and oleoylethanolamide (OEA) - were quantified in key regions of the descending pain-modulatory system and in brain areas associated with emotional aspects of pain processing. These included the prefrontal cortex (PFC), ipsilateral and contralateral amygdala, ipsilateral and contralateral hippocampus, PAG, RVM, and L4–L6 dorsal spinal cord, separated into ipsilateral and contralateral quadrats corresponding to the sciatic innervation territory. Expression of CB_1_, fatty acid amide hydrolase (FAAH) and monoacylglycerol lipase (MGL) was assessed in these regions. CB_2_ receptor expression was measured only in PFC and PAG due to lack of antibody availability for the other regions at the time of analysis. Plasma was obtained by centrifuging trunk blood at 3420 g for 15 minutes at 4 °C. All samples were snap-frozen and stored at −80 °C until analysis.

**Figure 1.**
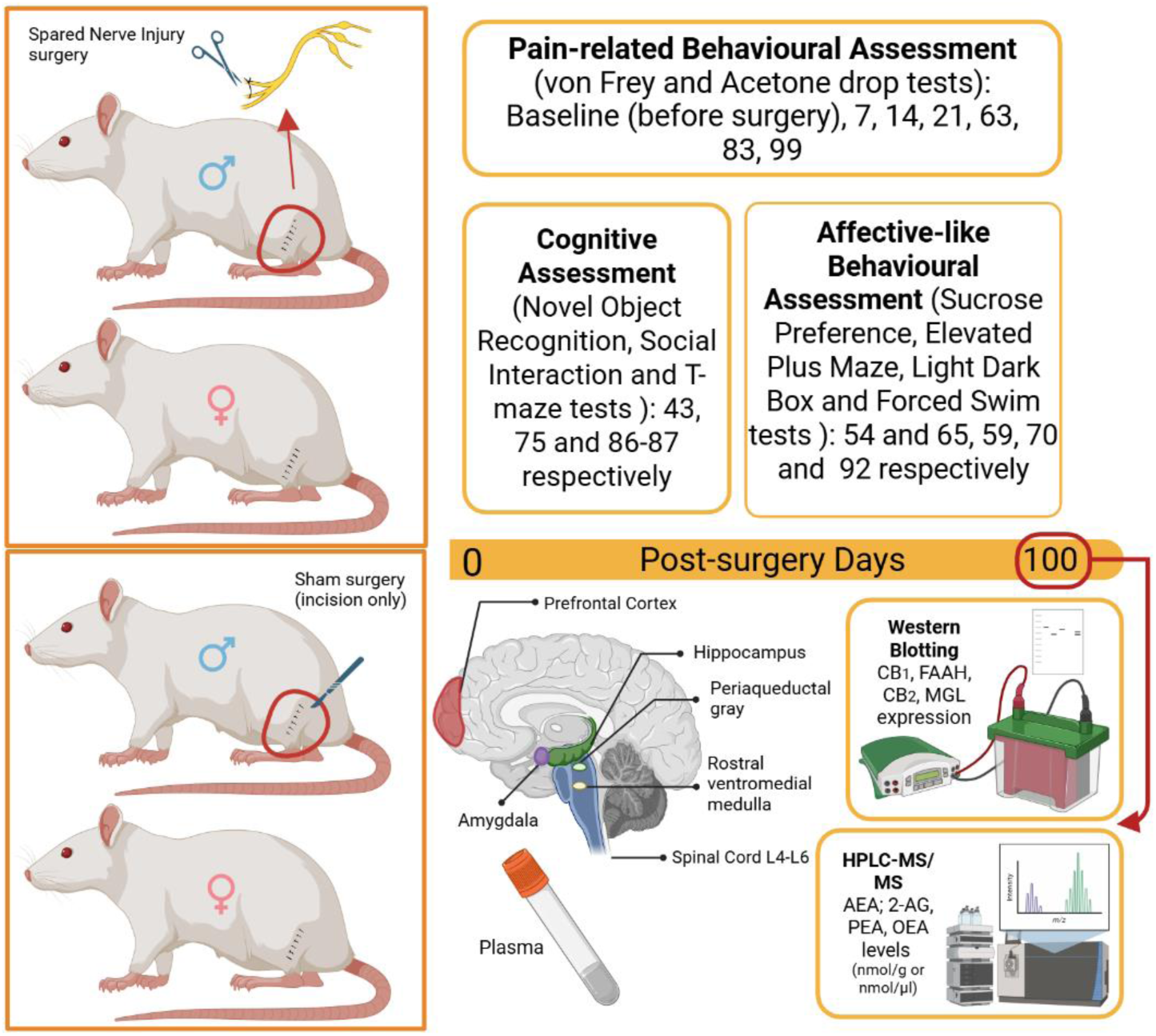
Schematic representation of the experimental design, and overview of the molecular and biochemical analyses performed.

### Quantification of endocannabinoid and related N-acylethanolamine (NAE) levels in brain, spinal cord and plasma samples using LC-MS/MS

Levels of endocannabinoids (AEA, 2-AG) and related NAEs (PEA, OEA) were measured as previously described by our group^2,4,11^. Brain and spinal cord tissues were homogenized in 500μl of 100% acetonitrile using an ultrasonic homogenizer for 6-10 seconds, followed by centrifugation at 16,060g for 15 minutes at 4°C. The supernatant (75μl) was transferred into a 1.5 ml microfuge tube, and 200μl of acetonitrile containing deuterated internal standards for cannabinoid ligands and related N-acylethanolamines (NAEs) were added (2.5ng d8-AEA, 50ng d8-2-AG, 2.5ng d4-PEA, 2.5ng d4-OEA). After vortex-mixing, 40μl of the solution was transferred to an HPLC vial for mass spectrometry analysis.

Similarly, plasma samples (200μl) were thawed on ice for 30 minutes, then centrifuged at 16,060g for 15 minutes at 4°C. The plasma was transferred to 1.5ml microfuge tubes, and 20μl of 100% acetonitrile containing the same concentration of internal standards described before, were added. Protein precipitation was induced by adding 1ml of 100% acetonitrile with 0.1% formic acid, followed by a 30-minute incubation on ice. After centrifugation, the supernatant was filtered using PTFE syringe filters. The eluate was dried, reconstituted in 100% acetonitrile, and transferred to HPLC vials for analysis.

A standard curve using a fourfold serial dilution was prepared by adding 25μl of 100% acetonitrile containing a known fixed amount of non-deuterated analytical standard (125ng 2-AG and 12.5ng AEA, PEA and OEA) to 75μl of 100% acetonitrile into the highest standard (Standard 10). 200μl of 100% acetonitrile containing known fixed amount of deuterated internal standard was added to each standard, using the same stock that was used in the samples. 40μl of each standard were added to HPLC vials.

HPLC analysis was performed using a Zorbax® SB C18 column with gradient elution (Mobile phases comprised of Solution A (HPLC-grade water with 0.1% formic acid(v/v)) and Solution B (100% acetonitrile with 0.1% formic acid (v/v)) with a flow rate of 0.2 mL/min). Reversed-phase gradient elution was initiated at 45% solution B for 1 minute, then ramped linearly up to 100% solution B for 4 minutes and held at 100% solution B until 12 minutes. At 12.1 minutes, when the assay run finished, the gradient returned to initial conditions for a further 4-5 minutes to re-equilibrate the column before the next injection. Analyte detection was carried out in electrospray-positive ionisation mode on an Agilent 1260 Infinity II HPLC system coupled to a SCIEX QTRAP 4500 mass spectrometer operated in triple quadrupole mode (SCIEX Ltd, Phoenix House Lakeside Drive Centre Park, United Kingdom). Analyte retention times for 2-AG, AEA, PEA and OEA were 8.2, 7.9, 8.4 and 8.5 minutes, respectively. Quantification was achieved by multiple reaction monitoring (MRM) mode, and analyte concentrations were determined by calculating the peak area ratio of each target analyte to its deuterated internal standard using Skyline Quantitative Analysis Software. The concentrations were determined by using a 10-point calibration curve constructed from a range of concentrations of the non-deuterated form of each analyte and a fixed amount of deuterated internal standard. The data were expressed in ng per g of tissue or 200μl of plasma, converted to nmol/g or μl, and further converted to pmol/g or μl if necessary.

### Western Blotting

Frozen samples were lysed in radio-immunoprecipitation assay (RIPA) buffer with protease inhibitor and non-ionic detergent n-dodecyl-β-D-maltoside (DDM) to preserve G-protein-coupled receptor integrity ^13,43,49^. After homogenization and incubation, centrifugation separated the supernatant, which was collected for analysis. Protein concentration was determined using a Bradford assay and adjusted to 30μg/15μl. Equalized protein samples underwent 8% SDS-PAGE without heating to avoid denaturation. Proteins were electroblotted onto nitrocellulose membranes and verified for transfer with Ponceau staining. Membranes were blocked and incubated overnight at 4°C with primary antibodies (against FAAH (1:150, α-rabbit, Cambridge Bioscience Cat# 101600), MGL (1:150, α-rabbit, Abcam Cat# Ab24701), CB_1_ (1:150, α-rabbit, Abcam Cat# Ab23703), CB_2_ (1:150, α-rabbit, Abcam Cat# Ab3561), and β-actin (1:1000, α-mouse, Sigma Aldrich Cat# 5441)). After washing, membranes were incubated with fluorophore-labelled secondary antibodies (1:10000, IRDye 800CW and IRDye 680LT, LI-COR Cat# 929-70020 and Cat# 929-71010). Blots were scanned with a LI-COR Odyssey imager and analysed using LI-COR Image Studio Lite to calculate protein intensity ratios, which were normalized to the control group (male-Sham). Membranes were stripped and re-probed with different target antibodies.

### Statistical analysis

Data were analysed using IBM SPSS Statistics version 26.0. Normality and homogeneity of variances were assessed with the Shapiro–Wilk test (p>0.05) and Levene’s test (p>0.05), respectively. A two-way ANOVA was used to analyse data containing two factors (sex and surgery), and a three-way ANOVA was applied to datasets including three factors (sex, surgery, and side: ipsilateral vs. contralateral). *Post hoc* comparisons were performed using the Student Newman Keuls (SNK) test. Associations between mechanical and cold hypersensitivity levels 24h prior euthanasia and molecular measures were evaluated using Spearman’s rank-order correlation analysis, appropriate for non-parametric data. Correlation coefficients (ρ) and corresponding p-values were reported. Data are presented as mean ± SEM. Graphical representations were generated using GraphPad Prism version 8.0.

## Results

### Overview of Pain-related Behavioural Characterization

As previously reported^3^, SNI induced robust and sustained hypersensitivity to mechanical and cold stimuli in both sexes, with females exhibiting greater hypersensitivity across all chronic time points. These established sex-dependent behavioural differences provided the foundation for the present study, which examined whether persistent neuropathic pain is accompanied by distinct alterations in ECS components within key pain-processing regions in males and females. To this end, we quantified endocannabinoid ligands and related enzymes/receptors across key nodes of the descending pain modulatory system, as well as limbic and cortical regions involved in emotional and cognitive aspects of pain. A summary of these findings is presented in Table 1.

**Table 1.** Summary of changes in components of the ECS following SNI, compared with Sham.

| Brain region | AEA | 2-AG | PEA | OEA | CB<sub>1</sub> | CB<sub>2</sub> | FAAH | MGL |
| --- | --- | --- | --- | --- | --- | --- | --- | --- |
| PFC | = | ↑M | ↑M | = | = | = | = | = |
| Hippocampus | = | M<br>*(I>C) | = | = | = |  | = | = |
| Amygdala | = | = | = | = | F *(I<C) |  | M and F<br>*(I<C) | = |
| PAG | = | = | = | = | = | = | = | ↑F |
| RVM | = | ↑M | = | = |  |  |  |  |
| Spinal cord | = | = | = | = | = |  | = | = |
| Plasma | = | = | = | = |  |  |  |  |

F, female; M, male; ↑, increased levels or expression; ↓, decreased levels or expression; =, no change.
(\*) Differences in ECS components between ipsilateral (I) and contralateral (C) sides of the same region.

### Prefrontal Cortex

Analysis of endocannabinoid levels and NAEs in the PFC (Figure 2A-D) showed a significant effect of surgery on 2-AG levels (F_1, 18_=5.361, p<0.05. *Post hoc* p<0.05), specifically a male-specific elevation of 2-AG levels following injury. Further analysis revealed a sex*surgery interaction effect on PEA levels (F_1, 20_=12.402, p=0.01. *Post hoc* p<0.05), specifically a significant elevation of this analyte in male-injured rats. The increase in 2-AG levels correlated with reduced mechanical hypersensitivity within male subjects, while the increase in PEA levels correlated with reduced mechanical and cold hypersensitivity (Figure 2E, F, G, H. Spearman’s correlation test: R_s_=0.8407, R_s_=0.8697, R_s_=0.8407, R_s_=-0.8986, p<0.05). Furthermore, elevated 2-AG levels in the PFC were significantly correlated with reduced anxiety-like behaviour, as indicated by increased exploration of open arms in the EPM test (Rs=0.93, *p*=0.022); however no significant correlations were found between 2-AG or PEA and cognitive or social measures in the T-maze or 3-chamber sociability test, respectively (supplementary material Figure G). None of these SNI-induced alterations in 2-AG and PEA levels were accompanied by concomitant changes in FAAH, CB_1,_ MGL or CB_2_ receptor expression in the PFC (Figure 2I-L and supplementary material Figure B).

**Figure 2.**
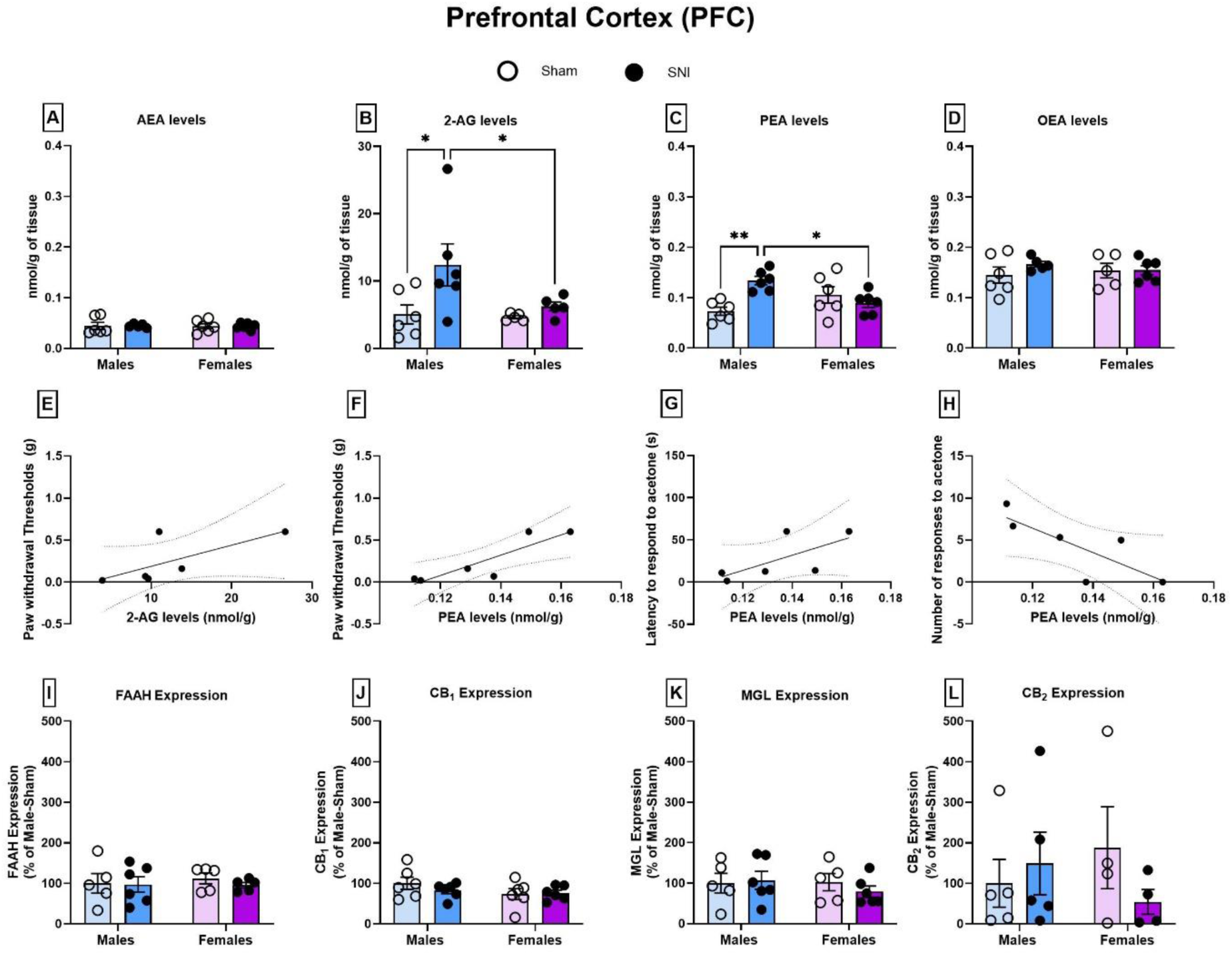
Levels of AEA, 2-AG, PEA, and OEA (A-D) and expression of FAAH, CB_1_, MGL, and CB_2_ (I-L) in the PFC of male and female rats. Nerve injury increased 2-AG and PEA levels in male rats only, an effect associated with attenuated mechanical (E, F) and cold (G, H) hypersensitivity. Data are expressed as mean ± SEM, n=5-6/group. Statistical significance: *p*<0.05. Correlation data points represent 6 individual XY pairs.

### Hippocampus

Analysis of endocannabinoid and NAE levels in the ipsilateral and contralateral hippocampus (Figure 3A–D) revealed a significant effect of surgery (F_1,37_=6.336, p<0.05), side (F_1,37_=37.047, p<0.05), and an overall of surgery*side interaction effect (F_1,37_=5.605, p<0.05) on 2-AG levels. *Post hoc* analysis confirmed reduced 2-AG levels in the contralateral hippocampus of male-Sham rats compared to the ipsilateral side. No significant effects of sex, side, or surgery were observed for AEA, PEA, or OEA.

**Figure 3.**
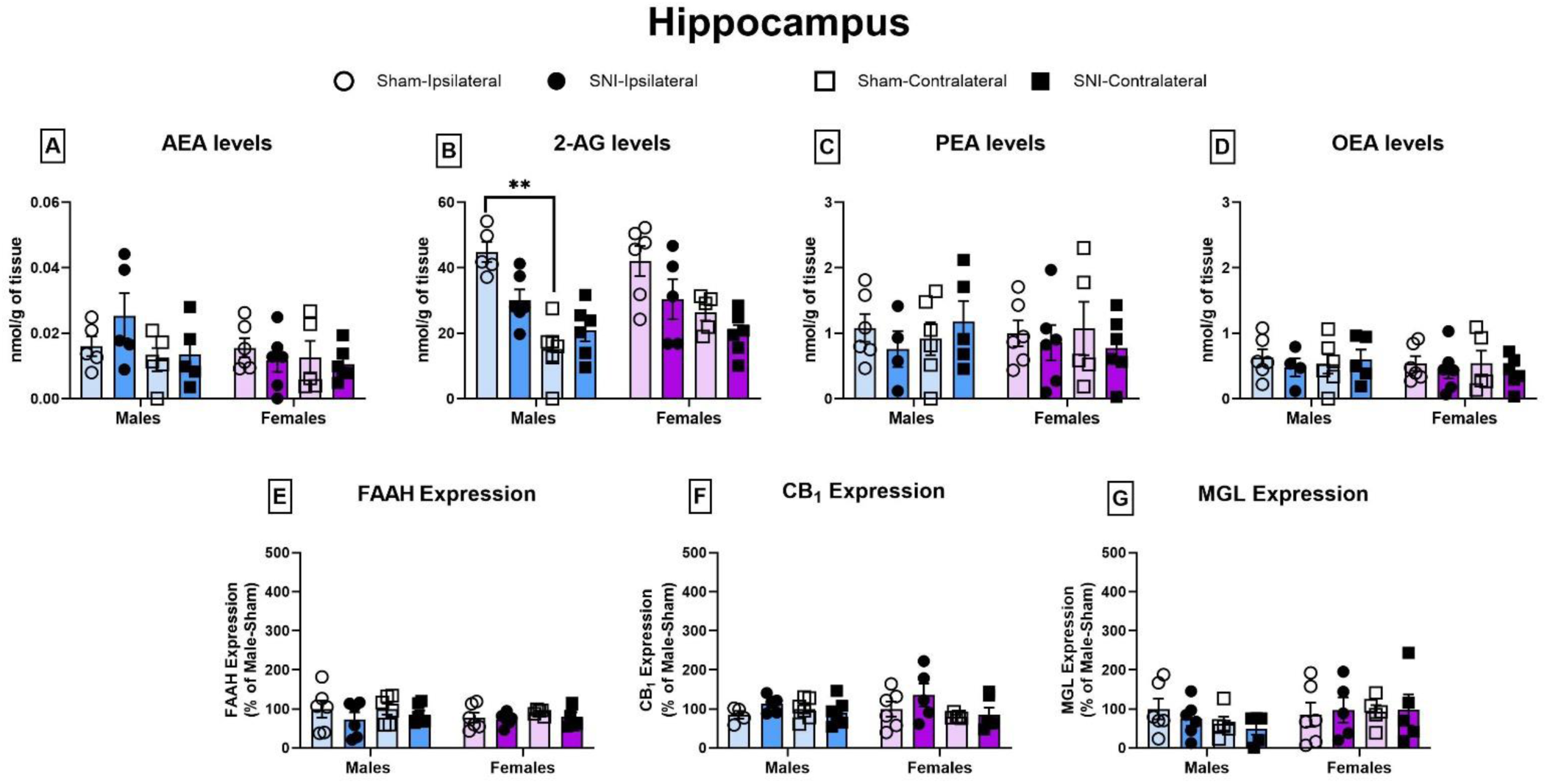
Levels of AEA, 2-AG, PEA, and OEA (A-D) and expression of FAAH, CB_1_, and MGL (E-G) in the hippocampus of male and female rats. Sham males showed lateralization of hippocampal 2-AG levels, with higher levels on the ipsilateral side. No injury-induced changes were observed in endocannabinoid or NAE levels, or in cannabinoid-related protein expression. Data are expressed as mean ± SEM, n=5-6/group. Statistical significance: *p*<0.05.

Analysis of FAAH, CB_1_ and MGL expression in the hippocampus revealed no significant differences across sex, surgery, or side (Figure 3E–G; Supplementary Figure C).

### Amygdala

Analysis of endocannabinoid and NAE levels in the ipsilateral and contralateral amygdala (Figure 4A-D) revealed a significant effect of side on levels of AEA (F_1,35_=6.387, p=0.016), PEA (F_1,34_ =5.635, p<0.05), and OEA (F_1,34_ =4.499, p<0.05), with higher levels in the contralateral side compared to the ipsilateral. However, subsequent *post hoc* pairwise comparisons did not reveal between-group differences. Similarly, the ANOVA showed a significant sex*surgery*side interaction effect for 2-AG levels (F_1,37_ =4.681, p<0.05), although *post hoc* analysis did not reveal any between-group differences.

**Figure 4.**
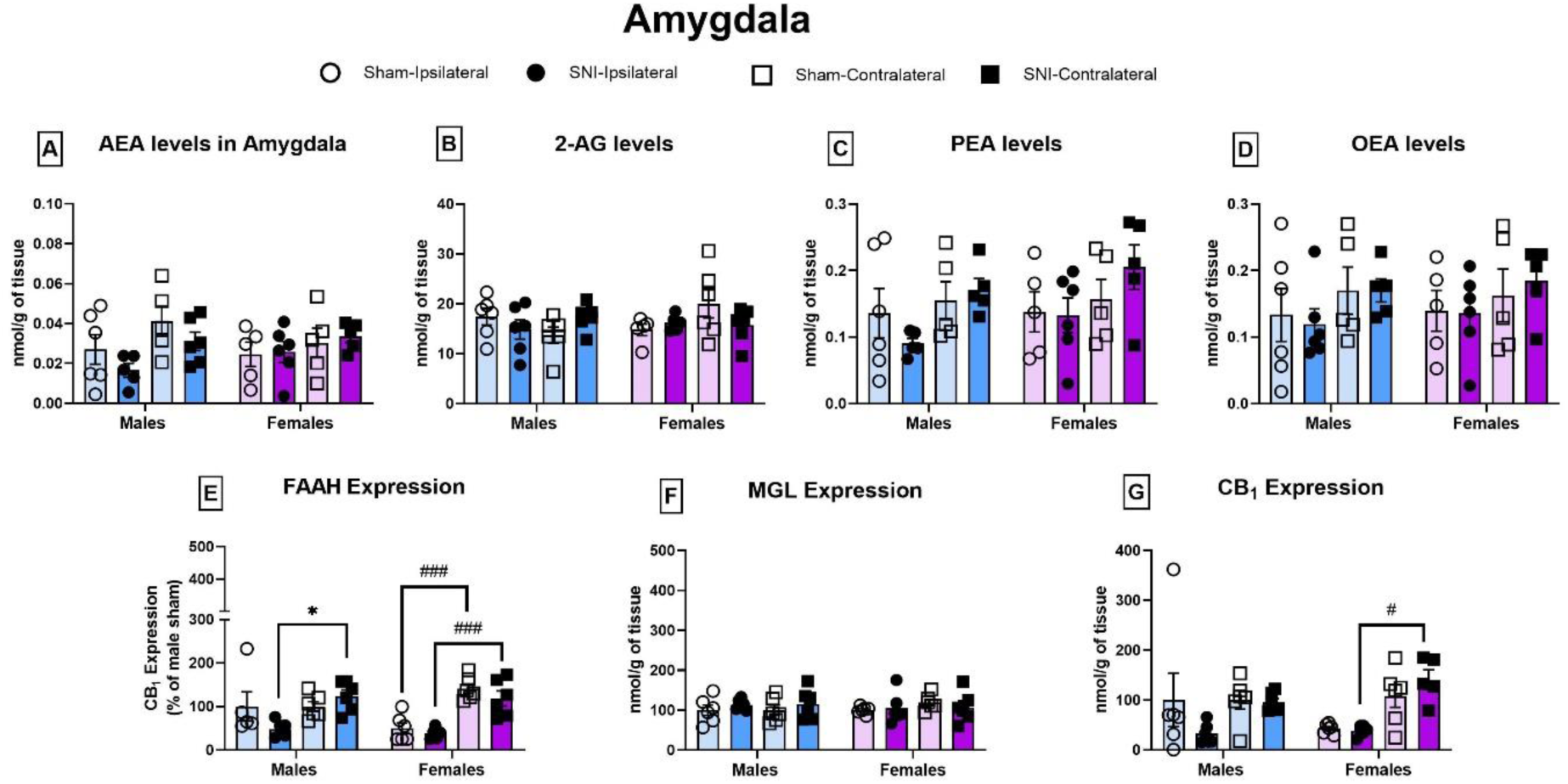
Levels of AEA, 2-AG, PEA, and OEA (A-D) and expression of FAAH, CB_1_, and MGL (E-G) in the amygdala of male and female rats. Nerve injury did not alter endocannabinoid or NAE levels, catabolic enzyme expression, or receptor density in either sex. However, higher FAAH expression was detected in the contralateral compared to the ipsilateral amygdala in females, a pattern also observed for CB_1_ expression in females. Data are expressed as mean ± SEM, n=6/group. Statistical significance: *p*<0.05.

Analysis of protein expression in the amygdala (Figure 4E-G; Supplementary Figure D) revealed an effect of side (F_1,38_=43.964, p<0.001) and sex*side interaction (F_1,38_=5.633, p<0.05) on FAAH levels. The subsequent *post hoc* tests confirmed higher FAAH expression in the contralateral side of the amygdala compared to ipsilateral, in male-SNI rats, and females regardless of surgery condition. Similarly, side differences were observed in CB_1_ receptor (F_1,39_=19.784, p<0.001), with higher expression in the contralateral amygdala than in the ipsilateral after SNI in females. No between-group differences were observed in MGL expression. These results suggest sex- and surgery-specific lateralization of endocannabinoid protein expression in the amygdala.

### Periaqueductal grey (PAG)

Analysis of endocannabinoid levels and NAEs in the PAG of male and female rats (Figure 5A-D) revealed no significant alterations following SNI surgery. There was a trend towards reduced 2-AG levels in female-SNI rats compared with female-Sham counterparts, but this effect failed to reach statistical significance.

**Figure 5.**
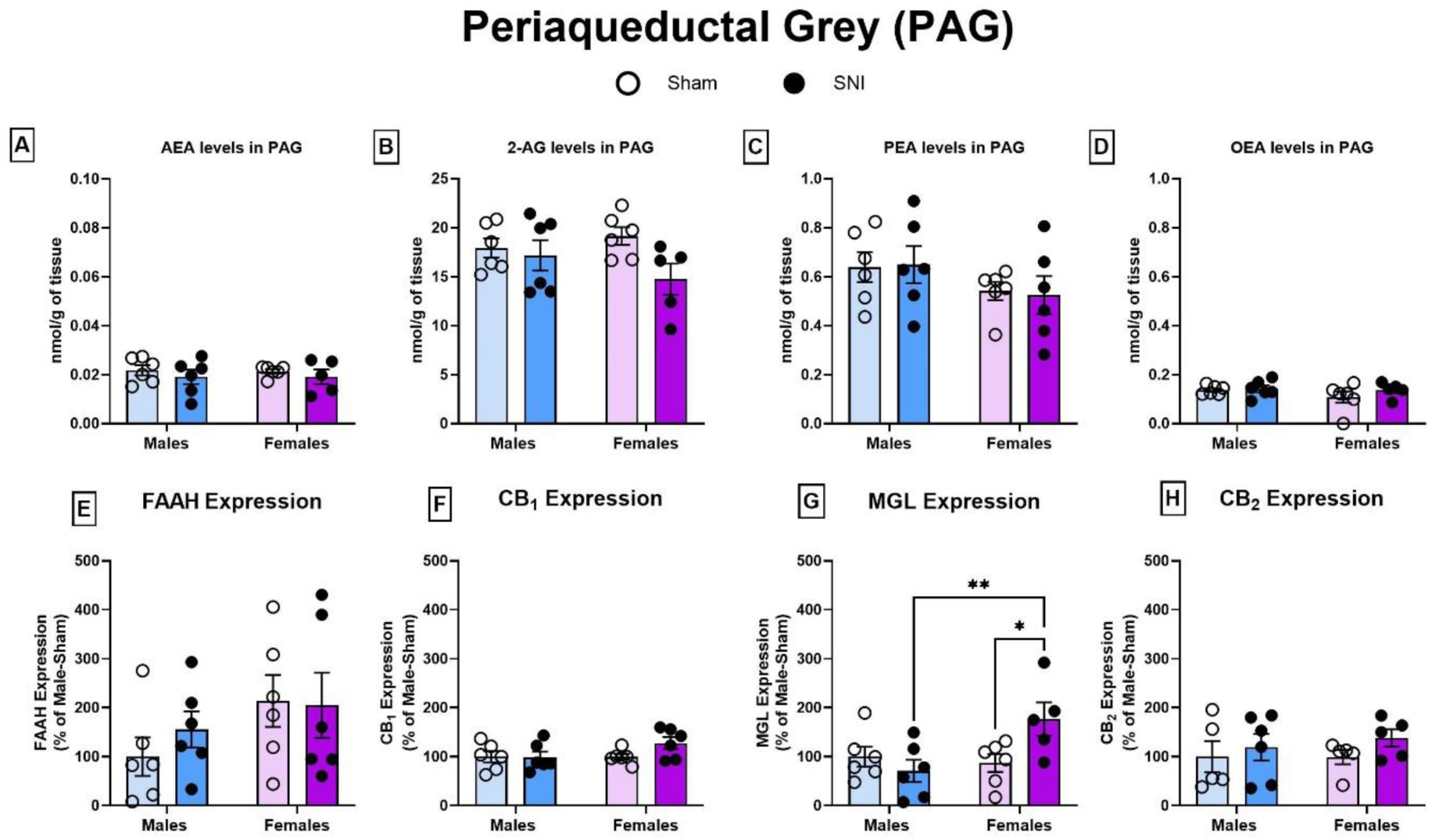
Levels of AEA, 2-AG, PEA, and OEA (A-D) and expression of FAAH, CB_1_, MGL, and CB_2_ (E-H) in the PAG of male and female rats. Although no significant changes were observed in endocannabinoid or NAE levels, a trend toward reduced 2-AG levels after nerve injury was detected in females. MGL expression increased in females following SNI compared to sex-matched controls, an effect associated with enhanced mechanical and cold sensitivity (I-J). Data are expressed as mean ± SEM, n=5-6/group. Statistical significance: *p*<0.05. Correlation data represents 10 individual XY pairs.

Analysis of ECS protein expression showed no differences in FAAH and CB_1_ (Figure 5E-H; Supplementary Figure E). However, there was a significant of sex*surgery interaction effect for MGL expression (F_1,19_=6.306, p<0.05), with *post hoc* analysis confirming increased levels in female-SNI rats compared to female-Sham controls. This MGL upregulation was in line with the trend toward reduced 2-AG levels in females after SNI.

### Rostral ventromedial medulla (RVM)

Analysis of endocannabinoids and NAEs in the RVM (Figure 6A-D) revealed an effect of surgery on 2-AG (F_1,19_=6.736, p<0.02) and OEA (F_1,16_=7.044, p<0.02) levels. Subsequent *post hoc* analyses confirmed an injury-specific increase in 2-AG levels in male rats.

**Figure 6.**
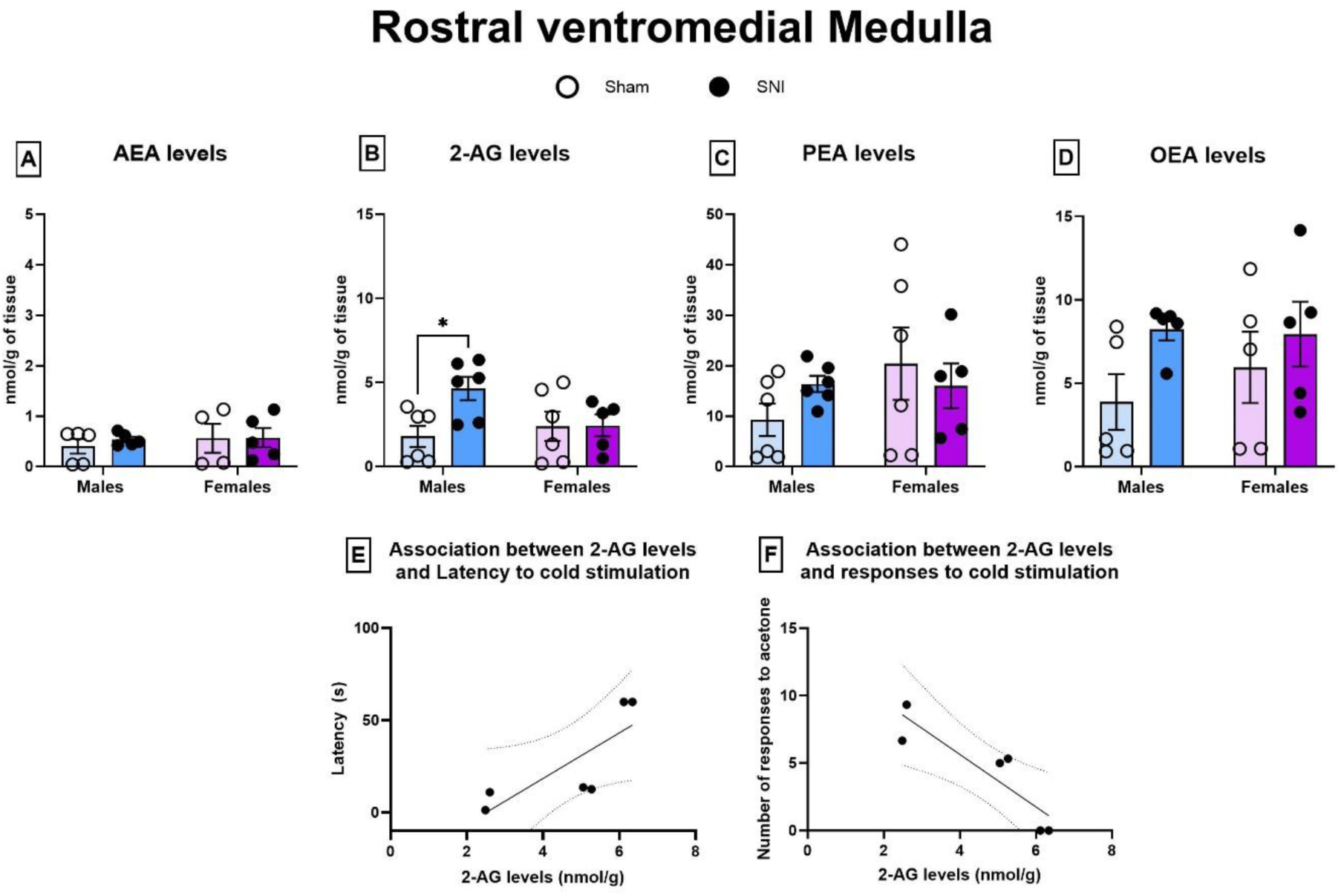
Levels of AEA, 2-AG, PEA, and OEA (A-D) in the RVM of male and female rats. Nerve injury increased 2-AG levels in males only, an effect associated with reduced cold hypersensitivity (E-F). Data are expressed as mean ± SEM, n=5-6/group. Statistical significance: *p*<0.05. Correlation data represents 10 individual XY pairs.

Correlation analyses revealed that increased 2-AG levels in male-SNI rats is associated with reduced mechanical (Rs=0.9276, p<0.02) and cold hypersensitivity (Rs=-0.8697, p<0.05) (Figure 6E-F).

### Spinal cord

Analysis of endocannabinoid and NAE levels in the ipsilateral and contralateral spinal cord (Figure 7A-D) revealed an effect of side in AEA (F_1,36_=7.311, p<0.01), 2-AG (F_1,33_=5.782, p<0.02) and PEA (F_1,36_=4.592, p<0.05) levels. However, further *post hoc* analyses confirmed no significant changes between ipsilateral and contralateral sides in either male or female rats.

**Figure 7.**
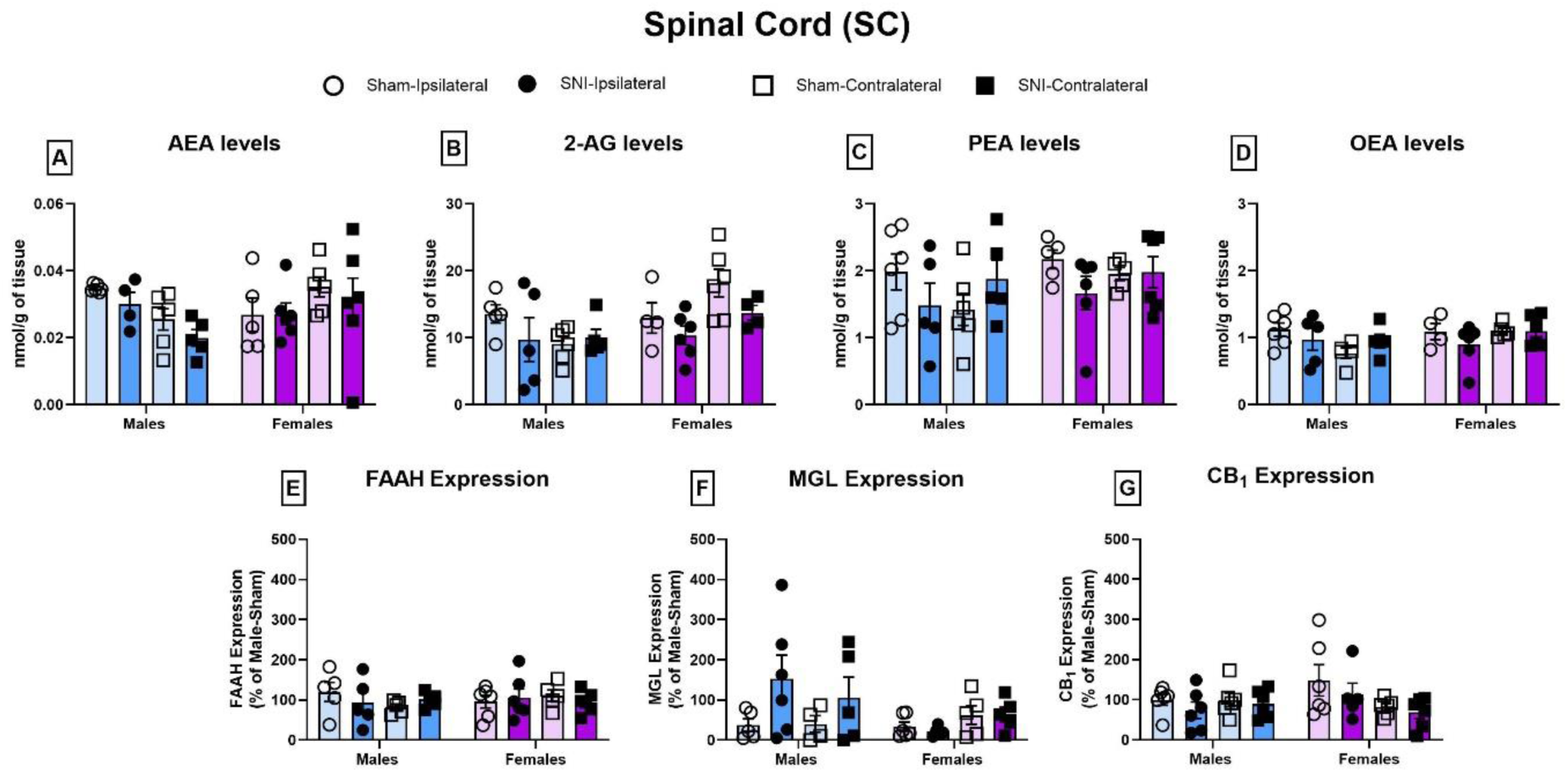
Levels of AEA, 2-AG, PEA, and OEA (A–D) and expression of FAAH, CB1, and MGL (E–G) in the spinal cord of male and female rats. Nerve injury did not alter any of the parameters investigated in either sex. Data are expressed as mean ± SEM, n=5–6/group.

Similarly, analysis of FAAH, CB_1_ and MGL expression (Figure 7E-G; Supplementary Figure F) revealed no significant differences among the groups and parameters analyzed.

### Circulating endocannabinoids and related NAE levels

Analysis of plasma endocannabinoid and NAE (Figure 8A-D) revealed no significant differences between sex or surgery on AEA, 2-AG, PEA or OEA levels.

**Figure 8.**
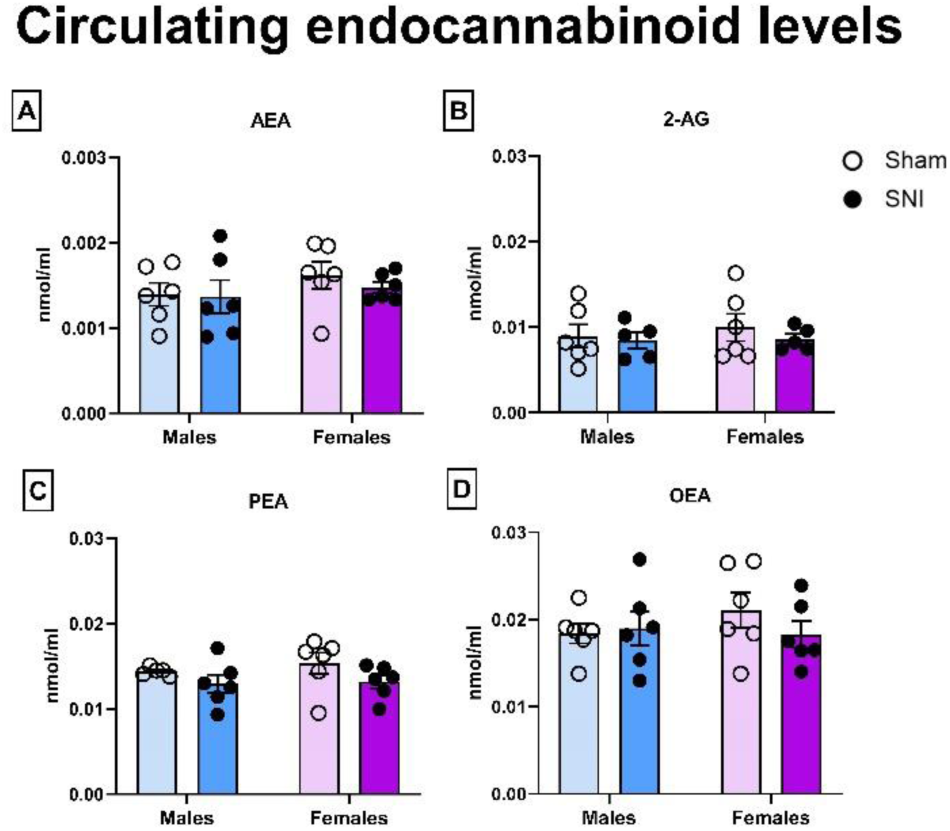
Circulating levels of AEA, 2-AG, PEA, and OEA (A-D) in male and female rats following Sham or SNI surgery. Data are expressed as mean ± SEM, n=6/group.

## Discussion

The present study demonstrates regionally distinct alterations in ECS signalling 100 days after SNI that differed between male and female rats. The principal injury-associated changes comprised increased 2-AG and PEA levels in the PFC and increased 2-AG in the RVM of males, together with increased MGL expression in the PAG of females. These findings identify persistent, region-dependent alterations in supraspinal ECS signalling during established neuropathic pain. In our previous studies, SNI produced persistent mechanical and cold hypersensitivity in both sexes, with greater hypersensitivity in females ^2,3^. These behavioural differences provided the framework for determining whether persistent neuropathic pain is accompanied by distinct patterns of ECS alteration across pain-processing regions in males and females.

In the PFC, SNI increased 2-AG levels in males, and exploratory within-group analysis showed that higher 2-AG levels were associated with reduced mechanical hypersensitivity. Endocannabinoid signalling within the PFC is increasingly recognized as an important regulator of persistent pain. Peripheral nerve injury can disrupt 2-AG-CB_1_ signalling in PFC circuits, altering the balance between inhibitory and excitatory transmission and contributing to abnormal cortical output ^26,35^. Against this background, the elevated PFC 2-AG levels observed at day 100 could be consistent with compensatory recruitment of endocannabinoid signalling during established neuropathic pain. However, because the present study assessed a single late time point and did not measure CB_1_ receptor function or synaptic signalling, the functional significance of this increase remains to be established.

SNI also increased PEA levels in the PFC of males, with higher PEA associated with reduced mechanical and cold hypersensitivity. PEA has previously been implicated in the modulation of neuropathic pain and associated cortical alterations. Chronic PEA administration restores glutamatergic synaptic homeostasis in the medial PFC and attenuates pain- and anxio-depressive-related behaviours following nerve injury ^19^. Together with these previous findings, the present association raises the possibility that increased PEA forms part of a compensatory response during persistent neuropathic pain. Importantly, neither the 2-AG nor PEA changes were accompanied by detectable alterations in CB_1_, CB_2_, FAAH or MGL expression in the PFC, suggesting that changes in ligand concentrations at this late stage are not reflected by corresponding changes in the abundance of the ECS proteins measured here. Determining the mechanisms responsible for these changes will require assessment of endocannabinoid synthesis, enzymatic activity and receptor function.

Exploratory analyses further identified a positive association between PFC levels of 2-AG and open-arm exploration in male SNI rats, whereas no significant associations were detected with cognitive or social measures. PEA levels were not significantly associated with these affective-cognitive outcomes. These observations should be interpreted cautiously because of the small number of animals included in the correlations and, importantly, because affective-cognitive behaviours were assessed substantially earlier than tissue collection. Accordingly, these associations do not establish a functional relationship between PFC ECS signalling and affective-cognitive outcomes but identify a potential relationship between PFC 2-AG and anxiety-related behaviour that warrants prospective investigation.

In contrast to the injury-associated changes observed in the PFC, SNI produced relatively limited alterations in ECS measures within the hippocampus and amygdala. No clear injury-related changes in hippocampal AEA, PEA or OEA levels or in FAAH, MGL or CB_1_ expression were detected. However, lateralization of hippocampal 2-AG levels was observed in males. Hemispheric specialization of hippocampal function has been described previously ^24,44^, although the functional significance of the 2-AG asymmetry observed here cannot be determined from the present data. Similarly, the amygdala exhibited lateralized differences in ECS protein expression, including FAAH and CB_1_, despite the absence of robust injury-related changes in endocannabinoid or *N*-acylethanolamine levels. Previous studies have demonstrated hemispheric specialization of amygdala function in nociceptive processing ^6,17,23^. Our findings extend this concept to components of the ECS and indicate that sex and hemisphere should be considered when examining ECS organization within limbic pain-processing circuits. However, the functional relationship between these lateralized molecular patterns and pain-related behaviour remains unresolved.

A distinct pattern emerged within the descending pain-modulatory system. The PAG, a major site of cannabinoid-mediated modulation of nociception ^16,22,28,34^, exhibited increased MGL expression following SNI in females. This was accompanied by a nonsignificant tendency toward lower 2-AG levels. As MGL is the principal enzyme responsible for 2-AG hydrolysis, this combination is consistent with altered regulation of 2-AG signalling in the female PAG after nerve injury. Pharmacological inhibition of MGL can enhance endocannabinoid signalling within the PAG and produce antinociceptive effects ^22,27^. One possibility, therefore, is that increased MGL expression could reduce the availability of 2-AG for CB_1_-mediated modulation of PAG circuits. Nevertheless, MGL activity, CB_1_ function and synaptic signalling were not directly assessed here, and the absence of a significant reduction in 2-AG concentrations means that this mechanistic interpretation requires direct experimental testing.

The PAG is also characterized by sex differences in the anatomical and functional organization of pain-modulatory pathways ^30–32^. In this context, the increase in PAG MGL observed in females identifies 2-AG metabolism as a candidate mechanism through which persistent nerve injury may differentially engage descending pain-modulatory circuitry in males and females. Importantly, this pattern contrasts with that observed in the RVM, where SNI increased 2-AG levels in males. Exploratory analysis further showed that higher RVM 2-AG was associated with reduced mechanical and cold hypersensitivity. Previous studies have demonstrated that CB_1_ receptor activation within the RVM modulates ON- and OFF-cell activity and produces antinociception ^36,46^, providing a plausible functional framework for this association. Together with the PFC findings, increased RVM 2-AG suggests that elevated 2-AG signalling accompanies the comparatively lower hypersensitivity observed in male SNI rats. Whether these changes represent compensatory antinociceptive responses requires direct manipulation of 2-AG signalling within these regions.

The contrasting PAG and RVM findings are particularly noteworthy because these structures form major components of the descending pain-modulatory system. Rather than indicating a uniform alteration of the ECS following nerve injury, the present results suggest that persistent neuropathic pain is associated with anatomically and sex-dependent patterns of ECS plasticity. In males, the predominant injury-associated changes involved increased ligand concentrations in the PFC and RVM, whereas in females the clearest alteration was increased expression of the 2-AG-catabolizing enzyme MGL in the PAG. These observations provide candidate molecular substrates for future studies examining how biological sex influences endocannabinoid regulation of descending pain control, but the present data do not establish that these molecular differences account for the behavioural differences between males and females.

No significant injury-related ECS alterations were detected in the L4-L6 spinal cord or plasma at day 100. These negative findings do not exclude functional ECS alterations at spinal or peripheral sites, particularly at earlier stages following injury or within specific cell populations that cannot be resolved using bulk-tissue analysis. Nevertheless, the absence of detectable changes in these compartments contrasts with the alterations observed in the PFC, PAG and RVM and highlights the regional specificity of the molecular phenotype observed at this late post-injury time point. Similarly, unchanged circulating AEA, 2-AG, PEA and OEA levels indicate that the central ECS alterations detected here were not accompanied by corresponding changes in plasma endocannabinoid or N-acylethanolamine concentrations, despite reports of altered peripheral ECS-related lipids in clinical neuropathic pain populations ^1,4^.

Several limitations should be considered when interpreting these findings. First, molecular analyses were conducted at a single late post-injury time point, preventing determination of the temporal sequence between ECS alterations and the development or maintenance of pain-related hypersensitivity. Second, the relatively small group sizes and exploratory within-group correlation analyses limit the precision of the observed associations. Third, measurements of endocannabinoid concentrations and protein expression in bulk tissue do not establish changes in enzymatic activity, receptor function, cellular localization or circuit-specific signalling. Accordingly, the present data cannot determine whether the observed ECS alterations causally modulate pain-related behaviour. Finally, ovarian-cycle stage was not incorporated into the experimental design, and future studies will be required to determine whether hormonal state contributes to variability in female ECS responses. Longitudinal and intervention-based studies targeting 2-AG signalling within the PFC, PAG and RVM will therefore be important for establishing the functional significance of the sex-associated patterns identified here.

## Conclusions and Future Directions

This study identifies regionally distinct patterns of ECS alteration in male and female rats during established neuropathic pain. In males, SNI was associated with increased 2-AG and PEA levels in the PFC and increased 2-AG in the RVM, with exploratory associations between these measures and reduced pain-related hypersensitivity. In females, SNI increased MGL expression in the PAG, identifying altered regulation of 2-AG metabolism within a key descending pain-modulatory region as a candidate mechanism for further investigation. In contrast, injury-related changes were not detected in the spinal cord or circulation at this late point, highlighting the regional specificity of the observed phenotype. Together, these findings demonstrate that biological sex is an important variable in chronic pain-associated ECS plasticity and identify PFC, PAG and RVM endocannabinoid signaling as targets for future causal investigation.

## Authors contributions

Alvaro Llorente-Berzal and David Finn contributed to conceptualization, revision and editing of the manuscript. Laura Boullon performed the experiments, the data collection and analysis, contributed to the conceptualization and wrote the original draft of the manuscript.

## Data availability

The data generated would be available upon request.

