## Supplementary Material for "Sex-dependent endocannabinoid adaptations in supraspinal pain-modulatory circuits after peripheral nerve injury"

In the following sequence of western blotting images, the protein samples were randomized across the blots ensuring representation of each sex and surgery groups (Figure A). The groups were: Male-sham, Female-Sham, Male-SNI and Female-SNI. A quality control (QC) sample was included in the last column of the gels. Samples were organised following the exact same scheme in all the brain regions analysed.

| Gel 1 | | |
|---|---|---|
| ID | Sex | Surgery |
| 1 | Male | SNI |
| 2 | Male | SHAM |
| 3 | Male | SNI |
| 11 | Female | SNI |
| 19 | Male | SHAM |
| 10 | Female | SHAM |
| 14 | Male | SNI |
| 4 | Female | SNI |
| QC | QC | QC |

| Gel 2 | | |
|---|---|---|
| ID | Sex | Surgery |
| 12 | Female | SHAM |
| 18 | Female | SNI |
| 13 | Male | SHAM |
| 8 | Male | SNI |
| 20 | Male | SHAM |
| 5 | Female | SNI |
| 24 | Female | SHAM |
| 7 | Male | SNI |
| QC | QC | QC |

| Gel 3 | | |
|---|---|---|
| ID | Sex | Surgery |
| 9 | Male | SNI |
| 23 | Female | SHAM |
| 17 | Female | SNI |
| 21 | Male | SHAM |
| 22 | Female | SHAM |
| 6 | Female | SHAM |
| 16 | Female | SNI |
| 15 | Male | SHAM |
| QC | QC | QC |

**Figure A**. Organization of samples position in the western blotting membranes generated for the data included in the manuscript. Samples labelled with ID 1, 12 and 9, from blots 1, 2 and 3 respectively, were the first samples loaded in the SDS-page gels after the chameleon duo pre-stained protein ladder (left side of the blot).

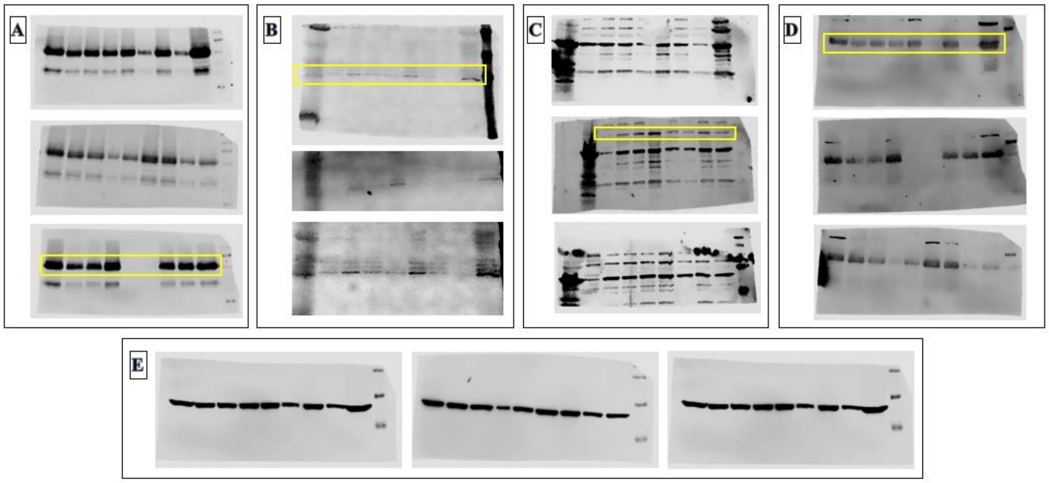


**Figure B**. Blots corresponding to (A) $CB_1$ (~53kDa), (B) $CB_2$ (~40KDa), (C) FAAH (~63KDa), (D) MGL (~35KDa) and (E) β-actin (~43kDa) expression in the PFC. The yellow rectangle indicates the band quantified to determine the expression of each protein.

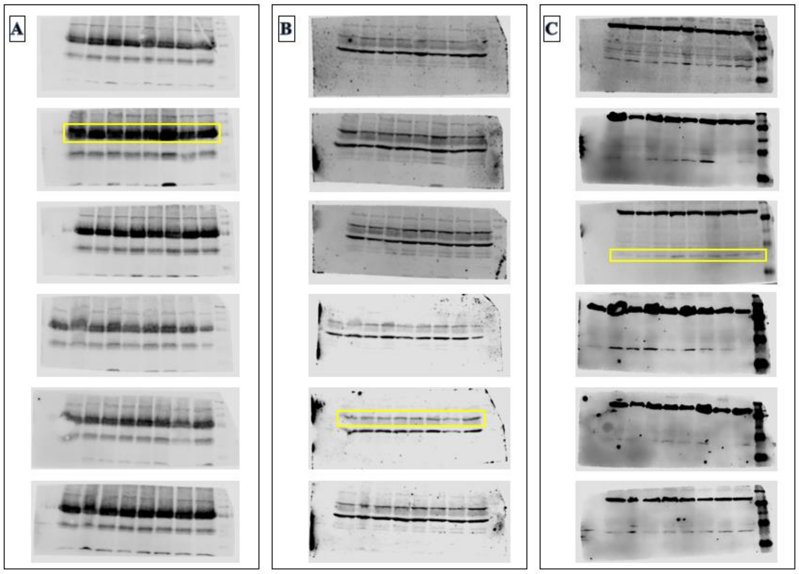


**Figure C**. Blots corresponding to (A) $CB_1$ (~53kDa), (B) FAAH (~63KDa) and (C) MGL (~35KDa) expression in the ipsilateral (first three images of each figure) and contralateral hippocampus (last three images of each group). The yellow rectangle indicates the band quantified to determine the expression of each target protein. Expression of β-actin was also quantified; however, images are not available.

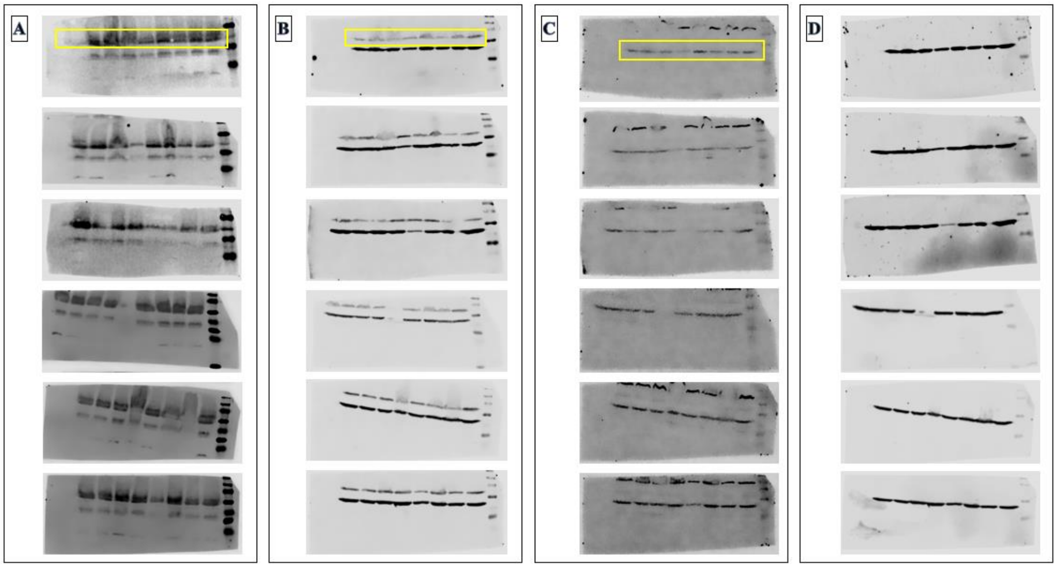


**Figure D**. Blots corresponding to (A) $CB_1$ (~53kDa), (B) FAAH (~63KDa), (C) MGL (~35KDa) and (D) β-actin (~43kDa) expression in the ipsilateral (first three images of each figure) and contralateral (last three images of each group) amygdala. The yellow rectangle indicates the band quantified to determine the expression of each target protein.

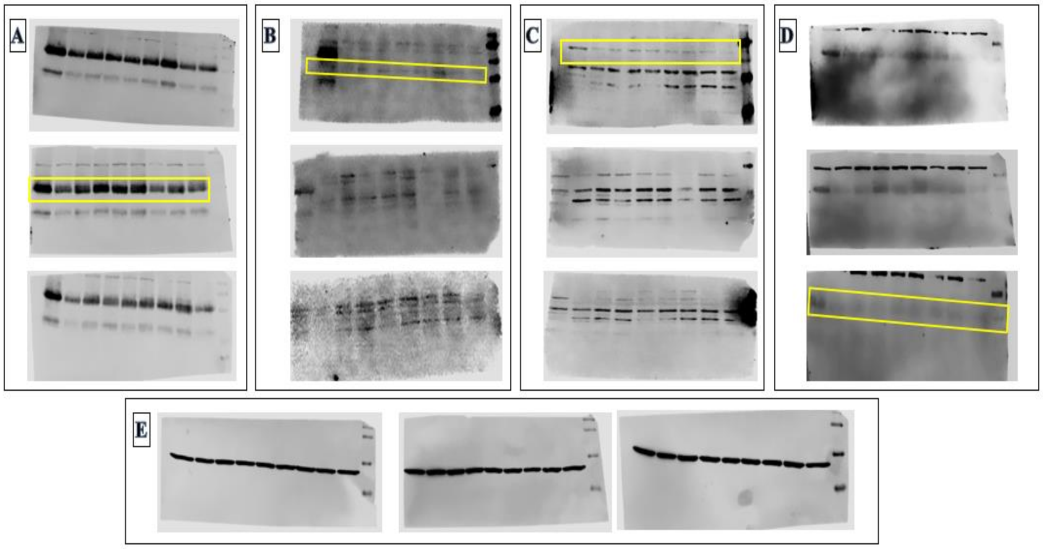


**Figure E**. Blots corresponding to (A) $CB_1$ (~53kDa), (B) $CB_2$ (~40KDa), (C) FAAH (~63KDa), (D) MGL (~35KDa) and (E) β-actin (~43kDa) expression in the PAG. The yellow rectangle indicates the band quantified to determine the expression of each target protein.

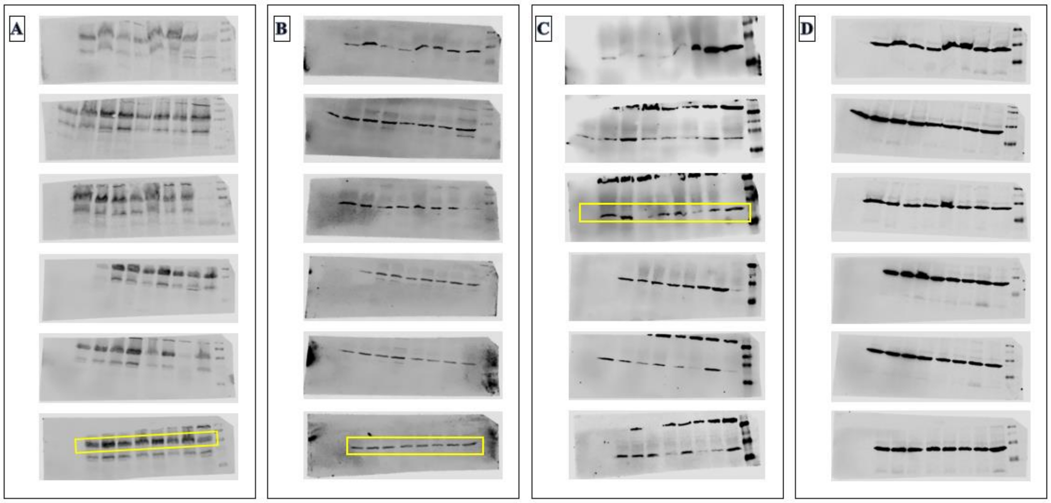


**Figure F**. Blots corresponding to (A) $CB_1$ (~53kDa), (B) FAAH (~63KDa), (C) MGL (~35KDa) and (D) β-actin (~43kDa) expression in the ipsilateral (first three images of each group: A, B, C, D) and contralateral (last three images of each group) spinal cord. The yellow rectangle indicates the band quantified to determine the expression of each target protein.

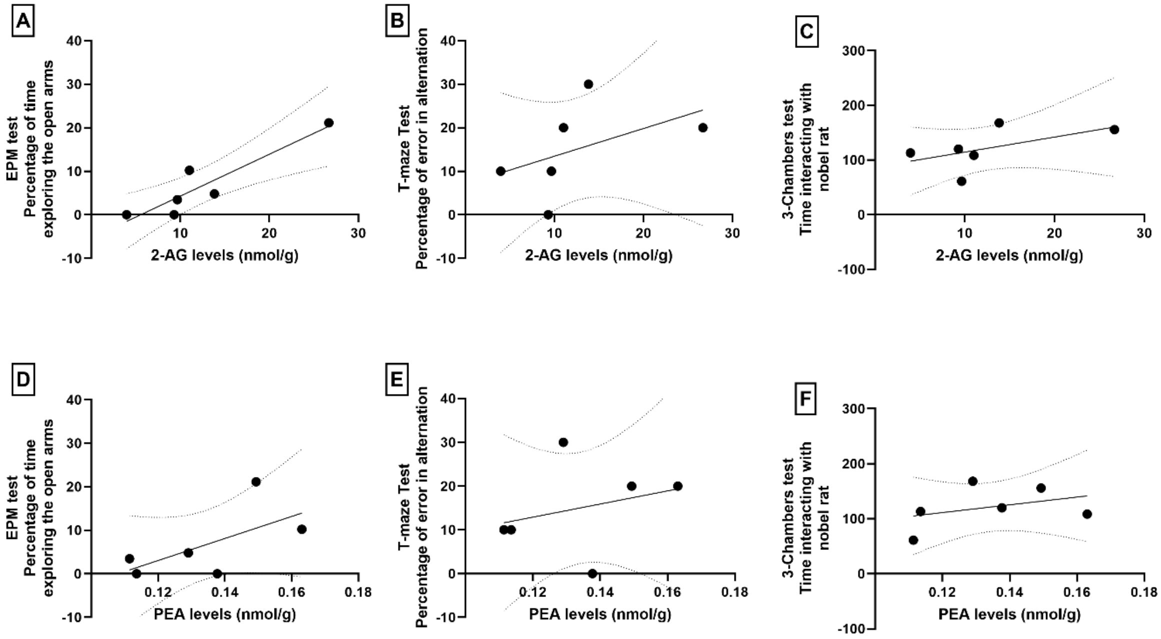


**Figure G**. Correlations between 2-AG (A-C) and PEA (D-F) levels and affective–cognitive behaviours in male SNI rats.

(A, D) Anxiety-like behaviour assessed in the Elevated Plus Maze (EPM); (B, E) Working memory in the T-maze; (C, F) Sociability in the Three-Chamber test.

Increased 2-AG levels in the PFC were positively correlated with the time spent exploring open arms in the EPM, indicating reduced anxiety-like behaviour within SNI males (Rs=0.93, $p$=0.022). No significant correlations were observed between 2-AG levels and cognitive (Rs = 0.79, $p$>0.05) or social behaviour (Rs=0.49, $p$>0.05). PEA levels were not significantly associated with any affective–cognitive measures after SNI (EPM: Rs=0.61; T-maze: Rs=0.32; Three-Chamber: Rs=0.26; all $p$>0.05). Correlation data points represent 6 individual XY pairs.
